# Bacterial composition and inferring function profiles in the biofloc system rearing *Litopenaeus vannamei* postlarvae at a low salinity

**DOI:** 10.1101/2022.01.01.474705

**Authors:** Hai-Hong Huang, Chao-Yun Li, Yan-Ju Lei, Wei-Qi Kuang, Wan-Sheng Zou, Pin-Hong Yang

**Author notes:** Correspondence, Hai-Hong Huang, Hunan University of Arts and Science. Changde, 415000, China. Co-first author.

## Abstract

This study aimed to investigate the bacterial composition and inferring function profiles in the biofloc system rearing *Litopenaeus vannamei* postlarvae (PL) at a low salinity condition. PL (~ stage 15) were stocked in four parallel tanks filled in water with a salinity of 5.0‰ at a density of 4000 individuals per m^3^ for a 28-days culture experiment, during which glucose was added as carbon source with a C:N of 20:1. At the end of experiment, water was sampled from each tank and pooled to extract microbial DNA for high-throughput sequencing of V3-V4 region of 16S rRNA gene. Results showed that the bacterial community at 28 d was dominated by phyla of *Proteobacteria* (45.8%), *Bacteroidetes* (21.1%), *Planctomycetes* (13.5%), *Chlamydiae* (10.3%) and *Firmicutes* (6.8%). A proportion of 81% inferring KEGG functions of this bacterial community associated with metabolism. Among functions relating to nitrogen metabolism, 48.5% were involved in the conversion of ammonia to glutamate, but the proportion of those involved in transformation among ammonia, nitrite and nitrate was 18.0% in total, inferring higher protein-synthesis but lower inorganic nitrogen-transformation capacities of the bacterial community. At the same time (28 d), high levels of total nitrogen (231.3±6.0 mg L^-1^) and biofloc (127.0±63.0 mL L^-1^), but low concentrations of ammonia (0.04±0.01 mg L^-1^), nitrite (0.2±0.1 mg L^-1^) and nitrate (12.9±2.5 mg L^-1^) were observed. The results supply a novel insight for understanding the function of bacterial community in the biofloc system nursing *L. vannamei* PL at a low salinity.

## 1 INTRODUCTION

*Litopenaeus vannamei* is the most important cultured crustaceans species in the world, whose culture production accounted for 52.9% of the total crustaceans production in 2018 (FAO, 2020). During culture of *L. vannamei*, after larvae period and before grow-out period, postlarvae (PL) are sequentially nursed for an intermediate period called the prenursery phase (Mishra, Samocha, Patnaik, Speed, Gandy, & Ali, 2008; Rezende Schleder, Silva, Henriques, Lorenzo, Seiffert, … & Vieira, 2018), to deliver large PL that are resistant to environmental conditions to increase the survival and growth performance. However, in the conventional prenursery system, due to intensively high stocking density (~ 60 PL L^-1^), controlling of toxic ammonia and nitrite is a big problem, as well as the biosecurity followed by operations such as water exchange for management of those nitrogen compounds (Samocha, 2010). Recently, biofloc technology (BFT) has been tried to be used to nurse *L. vannamei* PL (Khanjani, Sajjadi, Alizadeh, & Sourinejad, 2017; Rezende, Schleder, Silva, Henriques, Lorenzo, Seiffert, … & Vieira, 2018; Schveitzer, Lorenzo, Felipe, Pereira, Mourino, Seiffert, & Andreatta, 2017), due to the advantages of this technology on nitrogen assimilation *in situ* and pathogen control under minimal or zero water exchange conditions (Avnimelech, 2015; Huang, 2020). Besides, biofloc is also rich in nutrients, immunostimulants and bioactive compounds, such as essential amino acids (Li, Liu, Li, Deng, Abubakar, Lan, …, & Liu, 2018), unsaturated fatty acids (Ray, Leffler, & Browdy, 2019), lipopolysaccharide (LPS) and carotenoids (Ju, Ian, Lytha, Warren, Wenhaocedric, & Floyd, 2008; Vargas-Albores, Martínez-Córdova, Gollas-Galván, Garibay-Valdez, Emerenciano, Lago-Leston, … & Martínez-Porchas, 2019), contributing to growth-improvement, immune-enhancement and probiotic effects for shrimp (Panigrahi, Sundaram, Saranya, Kumar, Dayal, Saraswathy, & Gopal, 2019).

Biofloc is an aggregate of bacteria, protozoan, feces and organic detritus (Avnimelech, 2015; Schryver, Crab, Defoirdt, Boon, & Verstraete, 2008), among which the bacterial community is considered to play an of importance role for the advantages of biofloc mentioned above (Cardona, Gueguen, Magre, Lorgeoux, Piquemal, Pierrat, … & Saulnier, 2016; Ju, Ian, Lytha, Warren, Wenhaocedric, & Floyd, 2008). Previous studies show that the bacterial community in the marine water biofloc systems rearing *L. vannamei* is dominated by Proteobacteria, Bacteroidetes, Planctomycetes and Firmicutes (Huerta-Rabago, Martinez-Porchas, Miranda-Baeza, Nieves-Soto, Rivas-Vega, & Martinez-Cordova, 2019; Martínez-Córdova, Francisco, Estefanía, Ortíz-Estrada, Porchas-Cornejo, Asunción, & Marcel, 2018; Vargas-Albores, Martínez-Córdova, Gollas-Galván, Garibay-Valdez, Emerenciano, Lago-Leston, … & Martínez-Porchas, 2019; Xu, Xu, Huang, Hu, Xu, Su, … & Cao, 2019). Those phyla are important to the maintenance of a good water quality for biofloc system because many species belonging to them use organic matter and nitrogen compounds for growth, such as *Thiotrichaceae, Rhodobacteraceae* and *Saprospiraceae* (Cardona, Gueguen, Magre, Lorgeoux, Piquemal, Pierrat, … & Saulnier, 2016). Additionally, the inferring functions of bacterial community in biofloc system, including nitrogen metabolism, biosynthesis of nutrients, immunostimulants and bioactive compounds, indicate associations of bacterial community with shrimp growth and water quality (Hargreaves, 2013; Ju, Ian, Lytha, Warren, Wenhaocedric, & Floyd, 2008; Vargas-Albores, Martínez-Córdova, Gollas-Galván, Garibay-Valdez, Emerenciano, Lago-Leston, … & Martínez-Porchas, 2019). From this point of view, studies on bacterial composition and its inferring function profiles would supply deep insights on understanding the role of bacterial community in the biofloc system.

*L. vannamei* is an euryhaline species which could be cultured at low salinity conditions less than 1.0‰, which is a trend that will continue to grow globally (Roy, Davis, Saoud, Boyd, Pine, & Boyd, 2010). Osmoregulation capability of *L. vannamei* develops gradually in the postlarvae stages and can easily be acclimated to salinities as low as 0.5‰ by PL12 (Van Wyk, Davishodgkins, Laramore, Main, Mountain, & Scarpa, 1999), indicating that PL after stage 12 could be reared at a low salinity condition for a prenursery phase. And recently, in some studies, biofloc technology had been used for prenursery of *L. vannamei* PL under low salinity conditions (8-16‰) (Esparza-Leal, Amaral Xavier, & Wasielesky, 2016; Luo, Liu, Shan, & Tan, 2019). However, there is little information about bacterial community and its inferring functions in the low-salinity biofloc system until now.

This study aimed to investigate the bacterial composition and inferring function profiles in a biofloc system rearing *L. vannamei* PL at a salinity of 5.0‰, to deeply understand the function of bacterial community in the biofloc system.

## 2 MATERIAL AND METHODS

### 2.1 Ethic statement

The experiments were carried out at a local farm of Bifuteng eco-agriculture development Co., Ltd. (BEAD Co., Lat. 28°53’57.88” N, Long. 111°38’3.08” E) and Hunan University of Arts and Science (HUAS, Lat. 29°3’0.12” N, Long. 111°40’11.43” E), both of which locate in Changde, China, under principles in good laboratory animal care, according to the national standard of China (GB/T 35892-2018), ‘Laboratory animal-Guideline for ethical review of animal welfare’. The manuscript does not require ethical approval.

### 2.2 Preparation of culture water

Water with a salinity of 5.0‰ for culture experiment was prepared according to Ray and Lotz (2017), with some modifications. Briefly, artificial sea salt powder (Qianglong corporation, Tianjin, China), and food grade chemical reagents of KCl, MgCl_2_ and CaCl_2_ were added to tap water to make a final salinity of 4.96‰ (~5.0‰), with K^+^, Mg^2+^ and Ca^2+^ concentrations of 292, 934 and 318 mg L^-1^, respectively. Then, the water pH value was adjusted to near 8.0 by using food grade NaHCO3. After that, sterilization (10.0 mg L^-1^ chlorinedioxide) followed by neutralization (1.0 mg L^-1^ ascorbic acid) were executed according to the processes of previous studies (Gaona, Almeida, Viau, Poersch, & Wasielesky, 2017; Lara, Krummenauer, Abreu, Poersch, & Wasielesky, 2017).

### 2.3 Experimental design and operations

Four indoor tanks (width × length × depth = 2 × 2.5 × 1.3 m) of BEAD Co., each of which fixed five porous tubes (2.4 meter in length) in the bottom connecting with a 750-w whirl charging aerator (HG-750S, Sensen Group Co., Ltd., Zhoushan, China), were filled with 5.0 m^3^ culture water prepared above. *L. vannamei* PL (~PL15, 2.5±0.5 mg) which had been treated with a desalinating and acclimation procedure to adapt the experimental conditions (Luo, Liu, Shan, & Tan, 2019; Van Wyk, Davishodgkins, Laramore, Main, Mountain, & Scarpa, 1999), were kindly supplied from BEAD Co. and randomly assigned to the four experimental tanks with a stocking density of 4000 individuals per m^3^ for a 28-days culture. During the whole culture period, PL were fed with a commercial formulated shrimp diet (crude protein 40.0%, crude lipid 5.0%, crude fibber 5.0%, crude ash 15.0%, moisture 12.0%, Alpha corporation, Jiangmen, Guangdong, China), with a frequency of four times equally a day (6:00, 12:00, 18:00, 24:00), at feeding rates of 25-35% and 20-25% corresponding to the average shrimp weight of < 0.1 g and > 0.1 g, respectively, basing on the estimated total biomass and operations of Van Wyk, Davishodgkins, Laramore, Main, Mountain, & Scarpa (1999). Besides, glucose (food grade, carbohydrate content 90.0%, Fufeng biotechnology Co., Ltd., Hohhot, Inner Mongolia Autonomous, China) was added as exogenous carbon source, according to a carbon to an inputted nitrogen ratio (C:N) of 20:1 (Ebeling, Timmons, & Bisogni, 2006) contained in feed and carbon source inputted each time. The inputted C:N was the C:N contained in the inputted materials (feed and carbon source). Briefly, 0.9 g carbohydrate or 0.36 g carbon (40% carbon in carbohydrate) is contained in 1.0 g glucose with a carbohydrate content of 90.0%. Meanwhile, 0.4 g protein or 0.064 g nitrogen (16% nitrogen in protein) is contained in 1.0 g formulated feed due to the crude protein content of 40.0% (Avnimelech, 1999; Ebeling, Timmons, & Bisogni, 2006). Meanwhile, about 0.384 g carbon is contained in 1.0 g feed according to the calculating method of Kumar, Anand, De, Deo, Ghoshal, Sundaray, … & Lalitha (2017). Thus, 1.6 g glucose needed to be inputted when 1.0 g feed was fed to shrimp, according to the inputted carbon to nitrogen ratio (C:N) of 20:1 and the conception that 25% is theoretically converted as shrimp biomass and 75% would be lost to water body (Piedrahita, 2003). Throughout the whole experimental period, no water exchange was operated, but evaporating loss was complemented with dechlorinated tap water per week.

### 2.4 Water sampling and high-throughput sequencing of 16S rRNA gene

At 28 d, 50 ml water was collected from each tank and pooled as one sample according to the method by Martínez-Córdova, Francisco, Estefanía, Ortíz-Estrada, Porchas-Cornejo, Asunción, & Marcel (2018); and 200 ml sample was obtained in total. The water sample was filtered through a 0.22-μm pore size membrane. After that, the membrane was collected to extract bacterial DNA genome with an E.Z.N.A™ Mag-Bind Soil DNA Kit (OMEGA Bio-Tek, Inc., GA, USA), according to the manufacturers’ instructions. The genome was taken as template to amplify the V3-V4 region of 16S rRNA gene with the universal primers, 341F: 5’-CCTACGGGNGGCWGCAG-3’ and 805R: 5’-GACTACHVGGGTATCTAATCC-3’, in a 30-μl mixtures containing microbial DNA (10 ng/μl) 2 μl, forward primer (10 μM) 1 μl, reverse primer (10 μM) 1 μl, and 2X KAPA HiFi Hot Start Ready Mix 15 μl (TaKaRa Bio Inc., Japan), via a two-stage PCR procedure with a thermal instrument (Applied Biosystems 9700, USA): 1 cycle of denaturing at 95°C for 3 min, first 5 cycles of denaturing at 95°C for 30 s, annealing at 45°C for 30 s, elongation at 72°C for 30 s, then 20 cycles of denaturing at 95°C for 30 s, annealing at 55°C for 30 s, elongation at 72°C for 30 s and a final extension at 72°C for 5 min. Then, the PCR product was purified and recovered with MagicPure Size Selection DNA Beads (TransGen Biotech Co., Ltd., Beijing, China), and quantified and normalized with a Qubit ssDNA Assay Kit (Life Technologies, USA). After that, high-throughput sequencing was conducted on the Miseq platform (Illumina, USA) according to the standard procedure by Sangon Biotech (Shanghai) Co., Ltd.

### 2.5 Bacterial composition analysis

Analyses for the high-throughput sequencing data were carried out under the QIIME 2 (Quantitative Insights Into Microbial Ecology, Version 2019.10) framework (Bolyen, Rideout, Dillon, Bokulich, Abnet, Al-Ghalith, … & Caporaso, 2019). In brief, ambiguous nucleotides, adapter sequences and primers contained in reads, and short reads with length less than 30 bp were removed with the cutadapt plugin (Martin, 2011). After that, bases in the two ends of reads with quality score lower than 25 were trimmed. And then, reads were truncated to a same length from both ends equivalently. Reads with a length too low to be subjected to truncated operation were discarded. Thereafter, chimeras were filtered, and pair-ended reads were joined, dereplicated, and clustered to operational taxonomic units (OTU) with an identity of 0.97, by using the Vsearch tool (Rognes, Flouri, Nichols, Quince, & Mahé, 2016). then, Coverage, Chao1 index, Berger-Parker index, Shannon index and Simpson index, were computed with the diversity plugin of QIIME 2. Chao 1 is an index measure the theorical counts of OTUs in a sample, and represent the richness of OTUs; Berger-Parker and Simpson index are dominance or evenness indexes. Berger-Parker index expresses proportional importance of the most abundant species. Whereas, Simpson index represents probability of 2 individuals being conspecifics, and decreases with increase of dominance of predominant species. Shannon index is a synthetic index for judging the richness and evenness of a sample (Magurran, 2004). Finally, OTUs were annotated with the reference GreenGene database 13.8, collapsed at phylum, class, order, family and genus levels, respectively, and visualized with a multilayered pie chart produced by Krona (Ondov, Bergman, & Phillippy, 2011).

### 2.6 Deducing of inferring functions of bacterial community

Inferring functions of bacterial community were deduced with PICRUSt2 (Phylogenetic Investigation of Communities by Reconstruction of Unobserved States) (Douglas, Maffei, Zaneveld, Yurgel, Brown, Taylor, … & Langille, 2019). Briefly, OTUs were placed into the reference phylogeny of GreenGene database 13.8 (Barbera, Kozlov, Czech, Morel, Darriba, Flouri, & Stamatakis, 2018; Czech, Barbera, & Stamatakis, 2020). Then, hidden-state prediction was run to get 16S copy numbers of OTUs to normalize the predicted KEGG Orthology (KO) functions abundances (Louca & Doebeli, 2017). After that, KEGG pathways abundances were inferred based on predicted KO functions abundances (Ye & Doak, 2009). The results were visualized with a multi-layered pie chart produced by Krona (Ondov, Bergman, & Phillippy, 2011).

The interaction, reaction and relation network of KO functions involved in nitrogen metabolism was recolored in terms of their relative abundances with KEGG Mapper (Kanehisa & Sato, 2020), basing on the reference map of 00910 obtained from database of KEGG PATHWAY (https://www.kegg.jp/kegg/pathway.html).

In addition, Enzyme Commission (EC) functions abundances were also predicted and normalized based on 16S copy numbers obtained from the hidden-state prediction (Louca & Doebeli, 2017), to evaluate composition of inferring digestive enzymes (EC 3.1, EC 3.2 and EC 3.4). The results were visualized with a multi-layered pie chart produced by Krona (Ondov, Bergman, & Phillippy, 2011).

### 2.7 Analysis of contributions of bacteria on inferring functions

Contributions (relative proportions) of bacteria to inferring functions at different levels were analyzed with QIIME 2 (Version 2019.10) (Bolyen, Rideout, Dillon, Bokulich, Abnet, Al-Ghalith, … & Caporaso, 2019), respectively.

### 2.8 Zootechnical measurement

Thirty PL were selected randomly and individually weighed to the nearest 0.1 mg with an electric balance (AUX220, Shimadzu, Japan) each week. Weekly increment rate of body weight (wiR) and specific growth rate (SGR) were calculated according the following formulates.

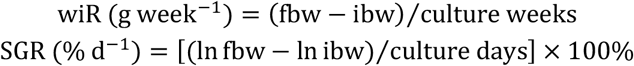

Wherein, fbw and ibw represented the final and initial mean body weight of PL, respectively. At 28 d when the experiment ended, all shrimp in each tank were harvested and counted individually to determine the survival rate (SR).

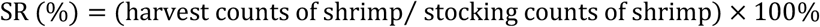

### 2.9 Water quality monitoring

Water temperature, dissolved oxygen and pH were detected each day by using electric analyzers (YSI-550A, Yellow Springs Instruments Inc., OH, USA). Water sample of each tank was passed thought 0.45-μm pore size microfilter (Xinya purification equipment Co., Ltd, Shanghai, China) once a week. Then, the filtrate was used to measure total ammonia nitrogen (TAN), nitrite, nitrate and carbonate alkalinity, and microfilter was used to measure total suspended solid (TSS), according to the standard methods (APHA, 1995). Water sample without filteration was used to determine total nitrogen (TN) weekly (APHA, 1995). Biofloc volume represented with settleable solids (SS) was determined per week with an Imhoff cone, by reading the sediment volume after a 15 min settlement of 1-liter water sample (Avnimelech, 2015).

### 2.10 Statistical analysis

Growth and water quality data in the present study were expressed as mean±standard deviation (SD) and statistically analyzed with the SPSS platform for windows (version 22.0, IBM Co., NY, USA). One-way ANOVA was executed as soon as normality distribution of data was proved with Shapiro-Wilk’s test. Tukey test or Dunnett’s T3 test were adopted for post hoc multiple comparisons of data with equal or unequal variances certified by using Levene’s test, respectively, if significant difference was found. Or else, non-parametric Kruskal-Wallis test was conducted, such as TAN, nitrite, biofloc volume (settleable solids) and carbonate alkalinity. Percentage data were submitted to arcsine transformation before statistical analyses. Differences were considered significant at *P* < 0.05.

## 3 RESULTS

### 3.1 Bacterial composition

The raw data produced from high-throughput sequencing has been deposited to NCBI Sequence Read Archive database with accession number of SRR12281666. After quality control, a total of 68411 reads with high quality was obtained and clustered to 3712 OTUs, with a Coverage of 0.96, Chao1 index of 13321.2, Berger-Parker index of 0.07, Shannon index of 7.6 and Simpson index of 0.98. The taxonomy profile was showed in Figure 1, where twenty-five phyla were assigned and dominated by Proteobacteria (45.8%), Bacteroidetes (21.1%), Planctomycetes (13.5%), Chlamydiae (10.3%) and Firmicutes (6.8%). At class level, Alphaproteobacteria (33%), Planctomycetia (11.8%), Saprospirae (11.4%), Chlamydiia (10.3%), Gammaproteobacteria (10.2%), Flavobacteriia (8%), Clostridia (3.6%) and Bacilli (3.2%), were the dominants (Figure 1). The first 10 predominant orders were Rhizobiales (14.1%), Rhodobacterales (13.0%), Saprospirales (11.4%), Chlamydiales (10.3%), Flavobacteriales (8.0%), Pirellulales (7.8%), Pseudomonadales (4.3%), Clostridiales (3.6%), Sphingomonadales (3.1%) and Lactobacillales (3.1%) (Figure 1). At other levels, there were nineteen families and eleven genera with proportion of more than 1.0% (Figure 1).

**FIGURE 1.**
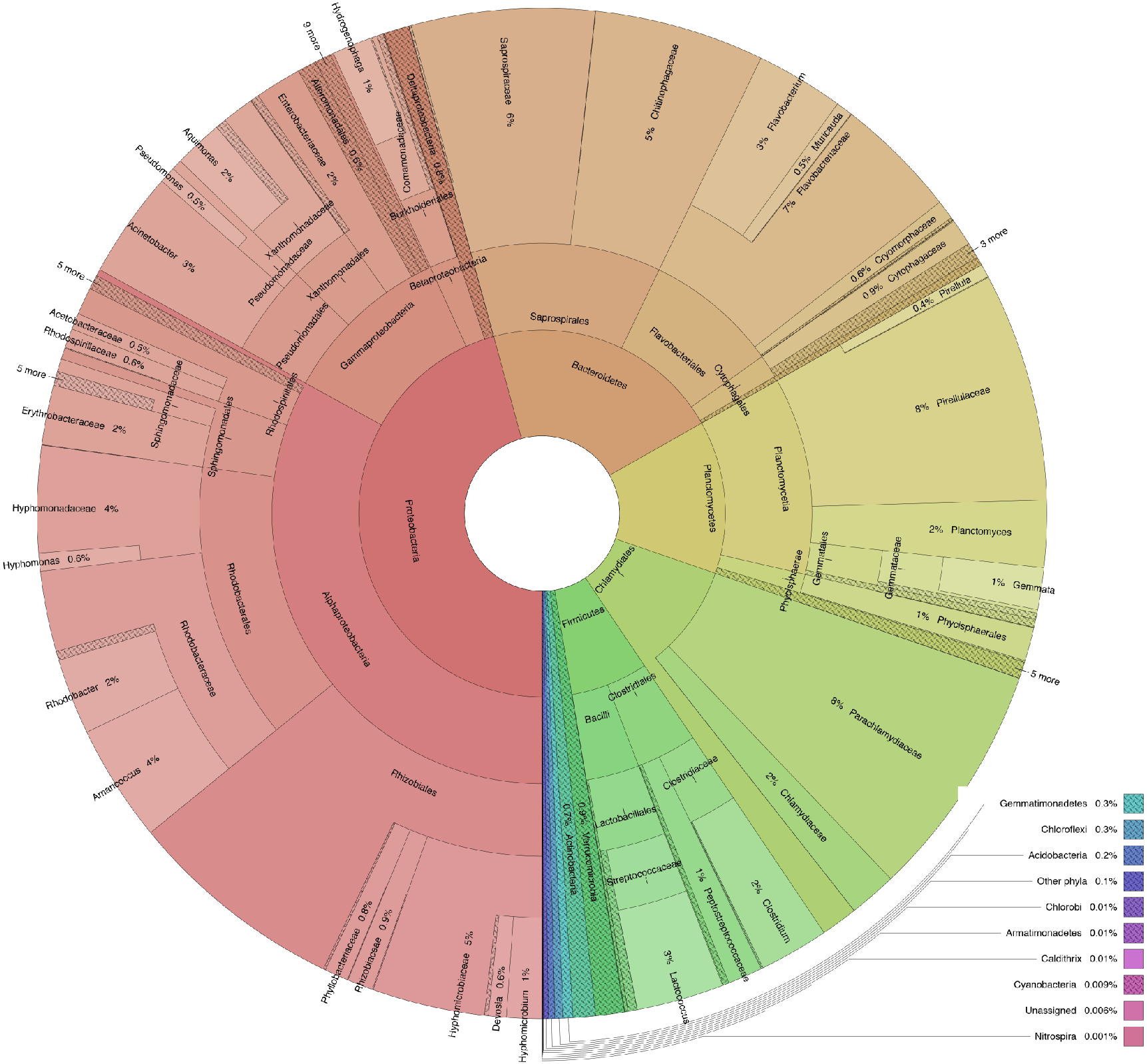
Five-layered pie chart for taxonomic compositions (relative abundance) of bacterial community at the end of the experiment (28 d) in the biofloc system rearing *Litopenaeus vannamei* postlarvae with a salinity of 5.0‰ at levels of phylum, class, order, family and genera.

### 3.2 Inferring functions of bacterial community

Inferring functions (KEGG pathways) profile of bacterial community at three levels (level 1-3) were analyzed. At level 1, Most inferring functions (81%) related to metabolism (Figure 2). The proportions of other level-1 functions, such as genetic information processing, cellular processes, environmental information processing and organismal systems were 11%, 4%, 2% and 0.4%, respectively (Figure 2). The most abundant level-2 functions were those relating to metabolism of nutrients, secondary metabolites and bioactive compounds, such as amino acids (21%, total of two categories, amino acid and other amino acids), carbohydrate (13%), lipid (8%), energy (5%), cofactors and vitamins (12%), terpenoids and polyketides (9%), xenobiotics (7%), and glycan (3%) (Figure 2). The most abundant Level-3 functions were Valine, leucine and isoleucine biosynthesis (amino acid metabolism), C5-branched dibasic acid metabolism (carbohydrate metabolism), biosynthesis of ansamycins and biosynthesis of vancomysin group antibiotics (metabolism of terpenoids and polyketides), fatty acid biosynthesis and synthesis and degradation of ketone bodies (lipid metabolism), and nitrogen metabolism (energy metabolism) (Figure 2).

**FIGURE 2.**
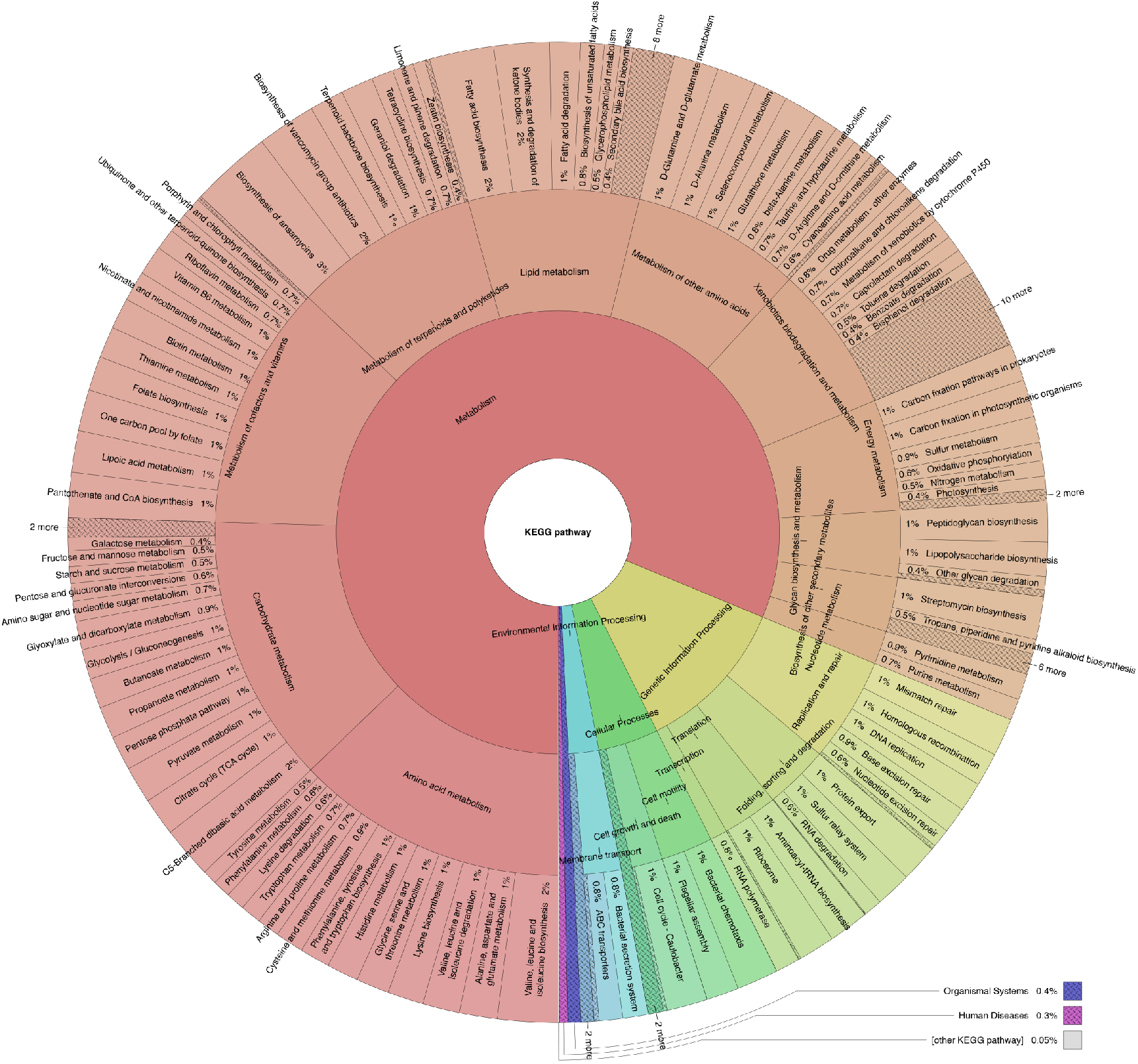
Three-layered pie chart for profiles (relative abundance) of level-1, level-2 and level-3 inferring functions of bacterial community at the end of the experiment (28 d) in the biofloc system rearing *Litopenaeus vannamei* postlarvae with a salinity of 5.()°oo. The inferring functions were represented with KEGG pathways.

Within level-3 KEGG pathway of nitrogen metabolism, the most important KO functions were referred to conversion of ammonia to glutamate which accounted for 48.5% (Figure 3). The proportion of KO functions relating to transformations among inorganic nitrogen compounds, such as nitrification, denitrification, and dissimilatory and assimilatory nitrate reduction, was 18.0% in total, including 1.2% for nitrite oxidization to nitrate, 7.0% for nitrate reduction to nitrite and 9.8% for nitrite reduction to ammonia (Figure 3).

**FIGURE 3.**
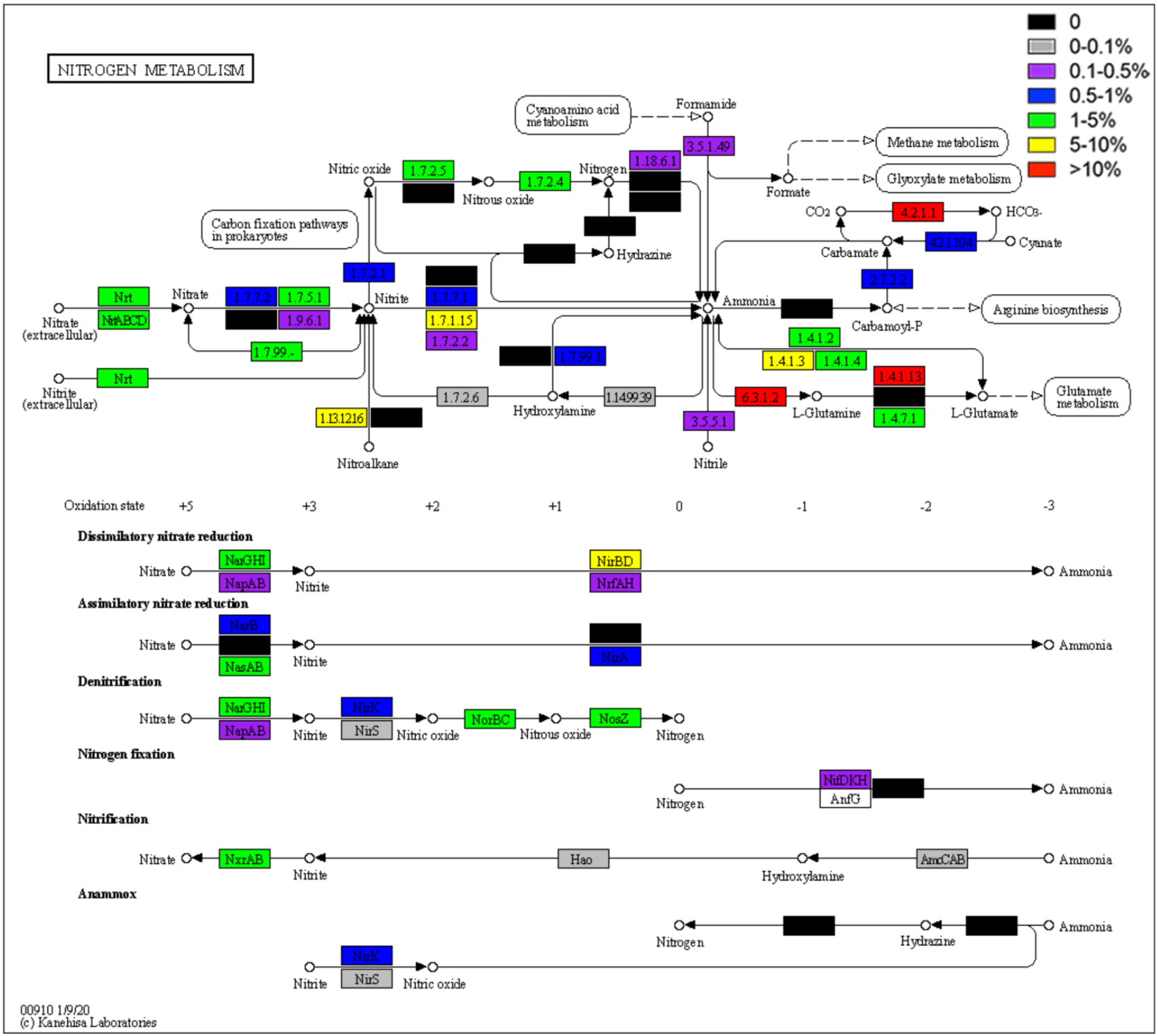
Profile of KEGG Orthology (KO) functions involved in the level-3 KEGG pathway of nitrogen metabolism of bacterial community at the end of the experiment (28 d) in the biofloc system w rearing *Litopenaeus vannamei* postlarvae ith a salinity of 5.0‰. Recolored in terms of relative abundances of KO functions, basing on the reference map of 00910 obtained from database of KEGG PATHWAY (https://www.kegg.jp/pathway/map00910).

Analysis results basing on predicted EC abundances showed that among enzymes with digestive activities, enzymes acting on ester bonds (EC 3.1), digesting carbohydrates (glycosylases, EC 3.2) and acting on peptide bonds (peptidases, EC 3.4) accounted for 53%, 15% and 32%, respectively (Figure S1). And in those three categories, the usual digestive enzymes (Fänge & Grove, 1979; Terra & Ferreira, 2012), such as triacylglycerol lipase, esterases (carboxylesterases), phosphatases; α-amylase, cellulase, chitinase, lysozyme, glucosidases; trypsin and aminopeptidases, were observed (Table 1).

**TABLE 1.**
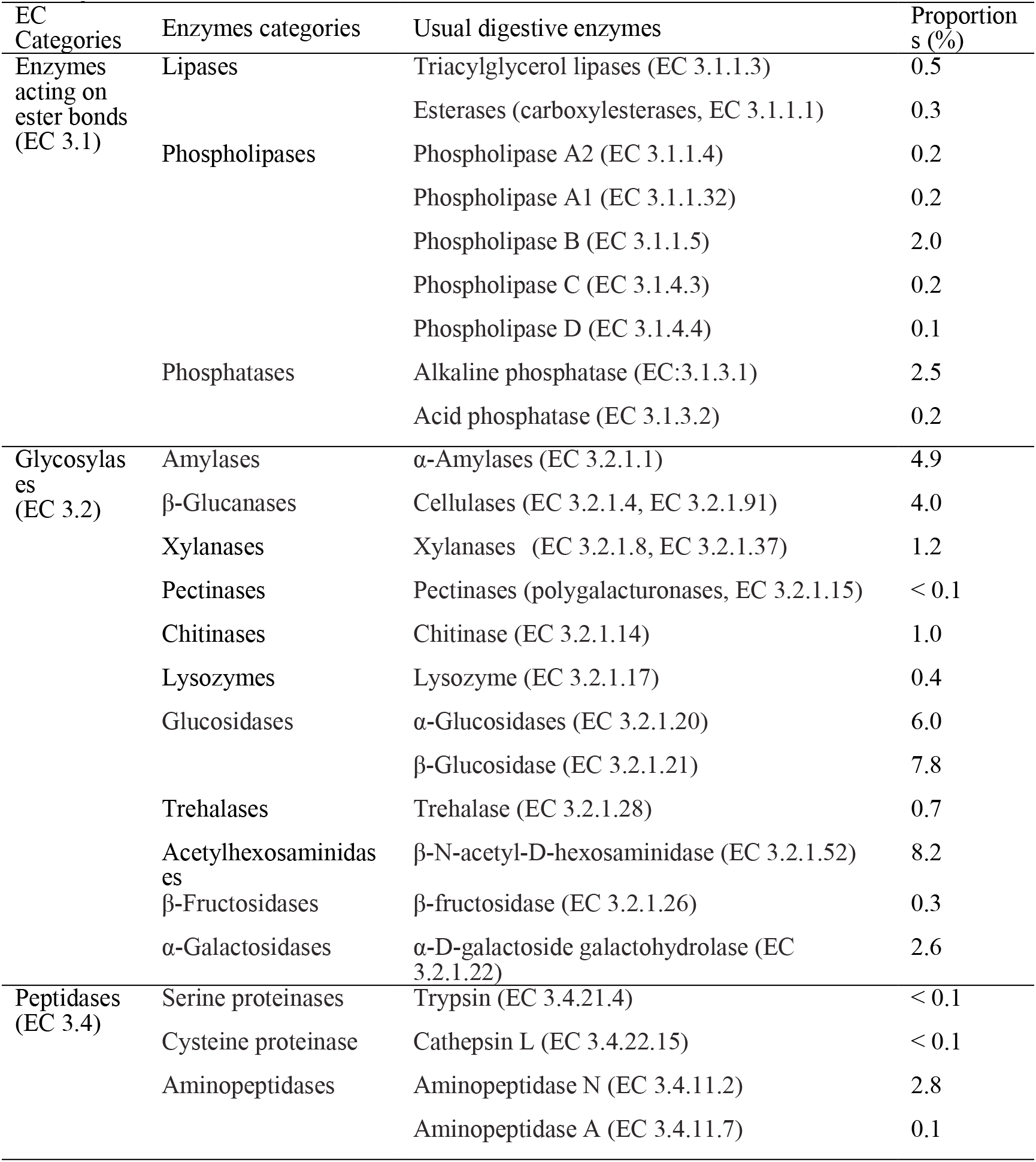
Relative proportions of predicted digestive enzymes in each EC categories of bacterial community at the end of the experiment (28 d) in the biofloc system rearing *Litopenaeus vannamei* with a salinity of 5.0%.

### 3.3 Contributions of bacteria on inferring functions

Four phyla Proteobacteria, Bacteroidetes, Planctomycetes and Firmicutes were found to be very important for the inferring functions (KEGG pathways), and contributed to approximate 89.4% of total functions (Figure 4). For level-1 functions, Proteobacteria, Planctomycetes and Firmicutes were most important phylum to metabolism, organismal systems and other functions, with contributions of 57.2%, 27.8% and 46.1%, respectively (Figure 4). Proteobacteria played very important roles almost to all level-2 and level-3 pathways relating to metabolism (Figures 5). KO functions contained in level-3 pathway of nitrogen metabolism were mainly contributed to phyla of Proteobacteria, Bacteroidetes, Planctomycetes and Firmicutes (Figure 6).

**FIGURE 4.**
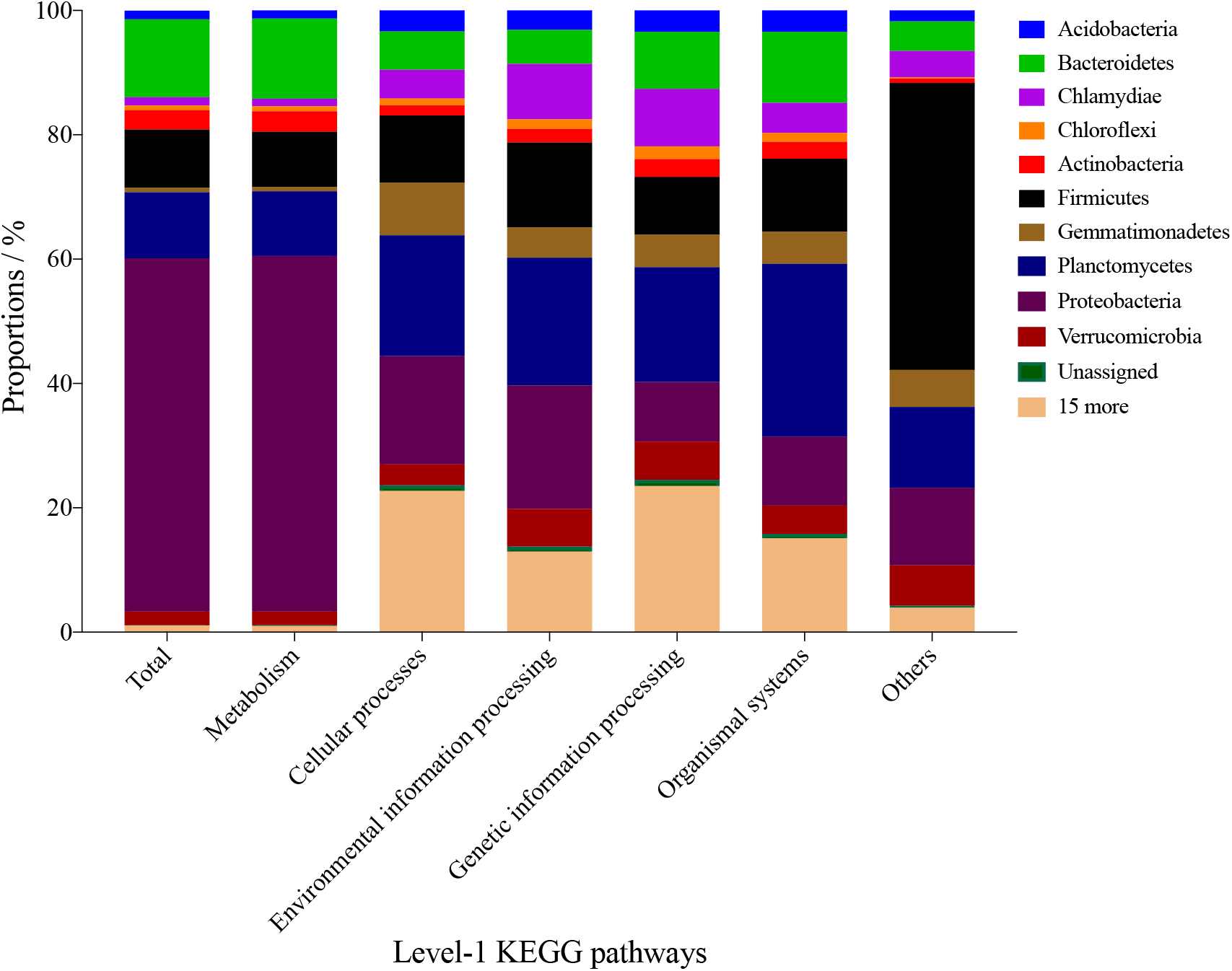
Stacked barchart displaying of contributions (relative proportions) of bacterial community at the end of the experiment (28 d) in the biofloc system rearing *Litopenaeus vannamei* postlarvae with a salinity of 5.0% at phylum level to level-1 inferring functions. The inferring functions were represented with KEGG pathways.

**FIGURE 5.**
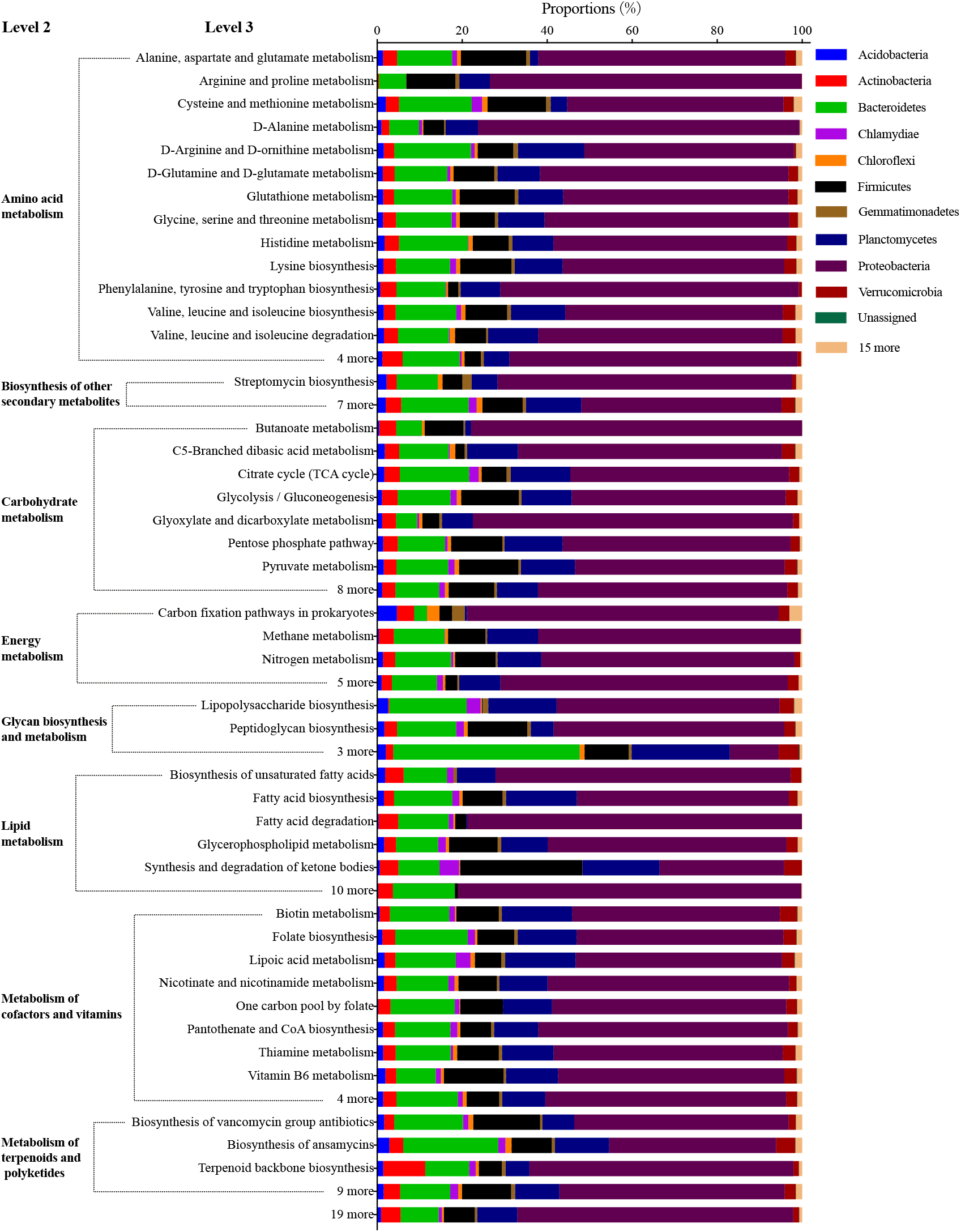
Stacked barchart of contributions (relative proportions) of phyla of bacterial community at the end of the experiment (28 d) in the biofloc system rearing *Litopenaeus vannamei* postlarvae with a salinity of 5.0‰ to level-3 KEGG pathways relating to metabolism (level 1).

**FIGURE 6.**
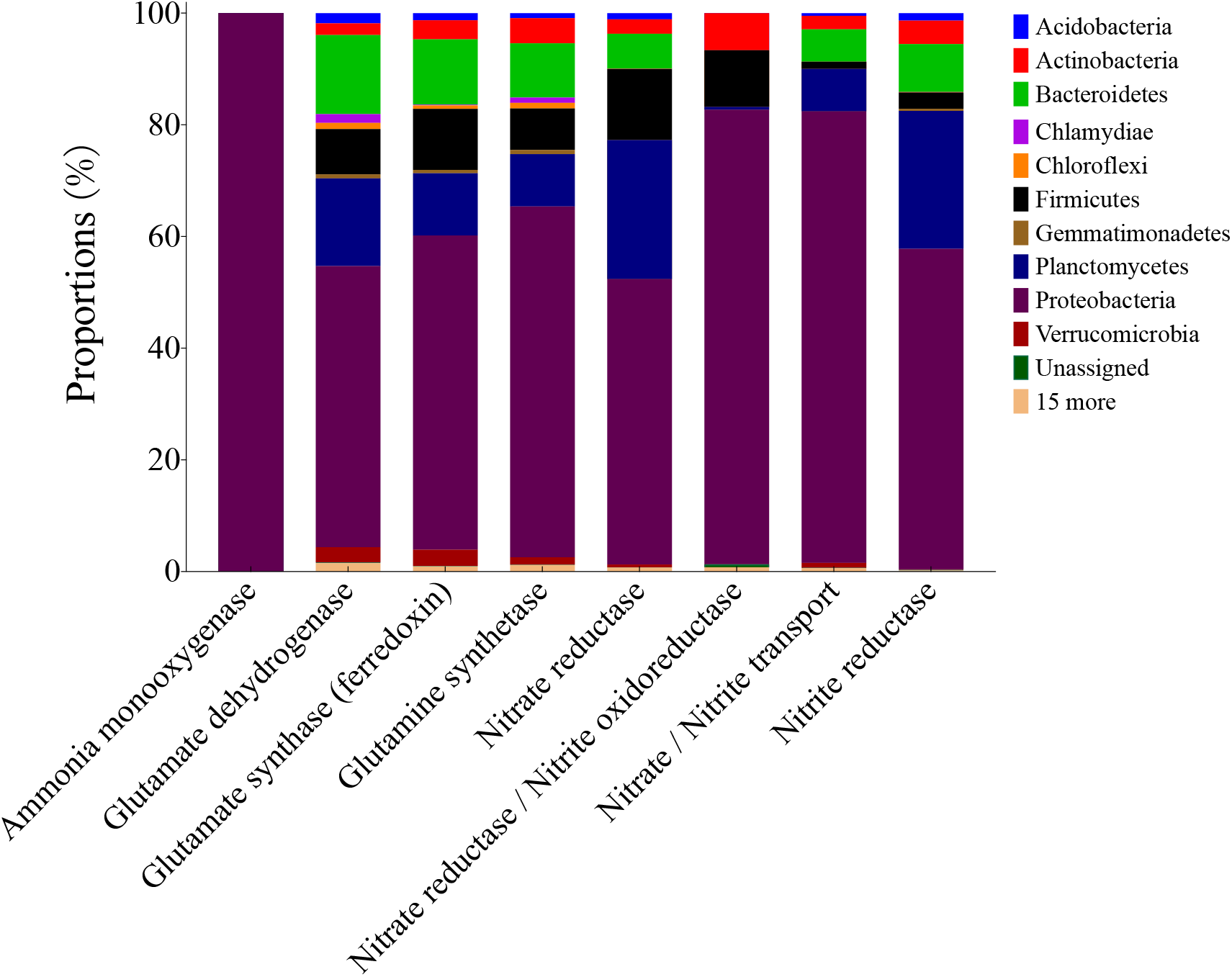
Stacked chart for contributions (relative proportions) of bacterial community at the end of the experiment (28 d) in the biofloc system rearing *Litopenaeus vannamei* postlarvae with a salinity of 5.0% at phylum level on the main KEGG Orthology (KO) functions relating to transformations among organic and inorganic nitrogen compounds contained in KEGG pathway of nitrogen metabolism (level 3).

### 3.4 Growth performance of PL and water quality

At harvest, 91.6±11.8% of PL was survived. The mean body weight of PL significantly increased to 120.9±16.7 mg at 28 d (*P* < 0.05, Figure S2). The concentrations of ammonia, nitrite and nitrate non-significantly oscillated during experiment (*P* > 0.05, Figure S3 a), with values of 0.04±0.01, 0.2±0.1 and 12.9±2.5 mg L^-1^ at 28 d, respectively. Whereas, the total nitrogen significantly accumulated to 231.3±6.0 mg L^-1^ at 28 d (*P* < 0.05, Figure S3 a). Similarly, biofloc and total suspended solids were also significantly accumulated to the levels of 127.0±63.0 mL L^-1^ and 240.0±62.0 mg L^-1^ at 28 d, respectively (*P* < 0.05, Figure S3 b). The value of pH displayed a significant decreasing trend (*P* < 0.05, Figure S3 c), with a final level of 7.4±0.1 at 28 d. The rest parameters, such as carbonate alkalinity, water temperature and dissolved oxygen were all stable over the whole experimental period (*P* > 0.05, Figure S3 c and d), with final values of 268.5±17.9 mg L^-1^ CaCO3, 29.8±0.3 ^o^C and 6.8±0.1 mg L^-1^ at 28 d, respectively.

## 4 DISCUSSION

### 4.1 Bacterial community properties

There are two key aspects about diversity of bacterial community, richness and evenness (Magurran, 2004). Chao 1 index measures richness based on rare species, increasing value means increasing of rare species, such as singletons. Several species dominance measurements, such as Berger-Parker index and Simpson index, are used to determine evenness. Berger-Parker index expresses proportional importance of most abundant species, increasing value means reduction in dominance of the most abundant species. Simpson index represents probability of 2 individuals being conspecifics, and increases with decreasing of dominance of the abundant species. Comprehensively, Shannon index indicates richness and evenness simultaneously. In the present study, high values of Chao1 index (13321.2), Shannon index (7.6) and Simpson index (0.98), but low value of Berger-Parker index (0.07) were observed, suggesting high diversity but low evenness of the bacterial community which was dominated by a few important species. In agreement with this, twenty-five phyla were observed in the present study, during which Proteobacteria, Bacteroidetes, Planctomycetes, Chlamydiae and Firmicutes, together represented 97.5% of the bacteria community. Similarly, in the biofloc systems rearing *L. vannamei* with marine water, 19-22 phyla were found, and Proteobacteria (26-73.88%), Bacteroidetes (6.57-85%), Planctomycetes (5.03-42%) and Firmicutes (6.48%) were also the most predominant ones (Huerta-Rabago, Martinez-Porchas, Miranda-Baeza, Nieves-Soto, Rivas-Vega, & Martinez-Cordova, 2019; Mariana, André, Felipe, Walter, Evelyne, Rafael, & Luciane, 2018; Martínez-Córdova, Francisco, Estefanía, Ortíz-Estrada, Porchas-Cornejo, Asunción, & Marcel, 2018; Vargas-Albores, Martínez-Córdova, Gollas-Galván, Garibay-Valdez, Emerenciano, Lago-Leston, … & Martínez-Porchas, 2019). Species belonging to those phyla, such as classes of Alphaproteobacteria, Planctomycetia, Saprospirae, Gammaproteobacteria and Flavobacteriia, and orders of Rhizobiales, Rhodobacterales, Saprospirales, Flavobacteriales, Pirellulales and Pseudomonadales, have strong adaptation capacity to different environments, especially conditions particularly rich in organic matter and suspended particles in the water column which are usually found in the BFT system, due to their abilities to use organic matter and nitrogen compounds for growth and particularity to attach to substrates for requirement of a support to grow on (Cardona, Gueguen, Magre, Lorgeoux, Piquemal, Pierrat, … & Saulnier, 2016; Kersters, Vos, Gillis, Swings, & Stackebrandt, 2006; Kirchman, 2002; Rank, 2009). Previous studies also revealed that in biofloc systems, although the proportions of the phyla mentioned above were low at initial, they should be predominant at last (Huerta-Rabago, Martinez-Porchas, Miranda-Baeza, Nieves-Soto, Rivas-Vega, & Martinez-Cordova, 2019; Martínez-Córdova, Francisco, Estefanía, Ortíz-Estrada, Porchas-Cornejo, Asunción, & Marcel, 2018; Xu, Xu, Huang, Hu, Xu, Su, … & Cao, 2019). Whereas, in the freshwater pond rearing fish with biofloc technology, Proteobacteria, Actinobacteria, Fusobateria, Chloroflexi and Saccharibacteria were the most dominant phyla (Liu, Li, Wei, Zhu, Han, Jin, … & Xie, 2019), different from the results observed in the present study with a low salinity condition. Whereas, the dominant phyla in the present study were found to be similar with those in the marine biofloc systems rearing *L. vannamei* (Huerta-Rabago, Martinez-Porchas, Miranda-Baeza, Nieves-Soto, Rivas-Vega, & Martinez-Cordova, 2019; Mariana, André, Felipe, Walter, Evelyne, Rafael, & Luciane, 2018; Martínez-Córdova, Francisco, Estefanía, Ortíz-Estrada, Porchas-Cornejo, Asunción, & Marcel, 2018; Vargas-Albores, Martínez-Córdova, Gollas-Galván, Garibay-Valdez, Emerenciano, Lago-Leston, … & Martínez-Porchas, 2019). It is hypothesized that the similar bacterial composition in the present study with those in seawater biofloc systems might be the result of successful adaptation of bacteria existing in marine-frying water to the low-salinity condition during the desalinating procedure and next operations. However, this adapting process is not clear and should be further studied for a deeper insight.

### 4.2 Properties of inferring functions of bacterial community

Vargas-Albores, Martínez-Córdova, Gollas-Galván, Garibay-Valdez, Emerenciano, Lago-Leston, … & Martínez-Porchas (2019) documented that most of the predicted level-1 KEGG pathways of bacterial communities in shrimp-culture marine water biofloc systems with amaranth and wheat as biofloc promoters associated with metabolism (50-53%), genetic information processing (19-21%), environmental information processing (12-15%), cellular processes and signaling (4-6%) and organismal systems (1-2%), in agreement with the results in the present study. However, compared to this previous study, the proportion of functions relating to metabolism is higher in the current study (81%), but those of genetic information processing (11%) and environmental information processing (2%) are lower. Different salinities between the present study and the previous study by Vargas-Albores, Martínez-Córdova, Gollas-Galván, Garibay-Valdez, Emerenciano, Lago-Leston, … & Martínez-Porchas (2019) might be partially responsible for those differences in inferring functions of bacterial community. Different environmental osmotic pressure under different salinity conditions would make bacteria adjust their physio- and biochemical behaviors. Additionally, other aspects, such as carbon source, C:N and age of shrimp, would also affect functions of bacterial community.

At the level 2, the most abundant functions were found to be associated to metabolism, such as metabolism of amino acids, carbohydrates, lipids, nucleotides, vitamins and cofactors, xenobiotics and other metabolites (Vargas-Albores, Martínez-Córdova, Gollas-Galván, Garibay-Valdez, Emerenciano, Lago-Leston, … & Martínez-Porchas, 2019), also similar with the results in the present study. The high abundance of those metabolic functions suggests that bacteria might be able to take advantage of the suspended substrate and suitable for degradation of organic matter (Vargas-Albores, Martínez-Córdova, Gollas-Galván, Garibay-Valdez, Emerenciano, Lago-Leston, … & Martínez-Porchas, 2019), and in turn to improve water quality and formation of bacterial biomass which could be taken as supplementary food for shrimp to improve growth performance.

### 4.3 Contribution profile of bacteria to inferring functions

In the present study, contributions of phyla Proteobacteria, Bacteroidetes, Planctomycetes and Firmicutes to inferring functions were ratable to their proportions in the total bacterial community, respectively. For example, the most abundant phylum *Proteobacteria* (45.8%) attributed to 56.8% of total inferring functions, and played very important roles on almost all function categories. Species belonging to this phylum, such as classes of Alphaproteobacteria and Gammaproteobacteria, orders of Rhizobiales, Rhodobacterales and Enterobacteriales, are widely dispersed in the environment and play important roles in the nutrient cycling and the utilization of organic compounds (Berman, 2012; Cardona, Gueguen, Magre, Lorgeoux, Piquemal, Pierrat, … & Saulnier, 2016). However, the attribution of Chlamydiae to functions was only 1.3%, although it was the fourth predominant phylum (10.3%). Vargas-Albores, Martínez-Córdova, Gollas-Galván, Garibay-Valdez, Emerenciano, Lago-Leston, … & Martínez-Porchas (2019) found that, compared to the taxonomic profile, the predicted functional profile shows a more effective pattern for representation of sample. This phenomenon should be deserved to note in future studies.

### 4.4 Inferring influence of bacterial community on growth performance of PL

Previous studies showed that formation of biofloc was found to be favorable to the growth of PL (Kuhn, Boardman, Lawrence, Marsh, & Flick, 2009; Xu & Pan, 2012). Generally, biofloc is considered to be a complementary food to improve growth of shrimp, due to rich in protein, lipid, amino acid and other bioactive compounds (Ju, Ian, Lytha, Warren, Wenhaocedric, & Floyd, 2008; Kuhn, Boardman, Lawrence, Marsh, & Flick, 2009; Xu & Pan, 2012). In the present study, inferring functions of bacterial community relating to metabolism of nutrients were also found, such as biosynthesis of essential and non-essential amino acids, unsaturated fatty acids, and cofactors and vitamins. Moreover, inferring digestive enzymes were also observed, which might be helpful for growth of PL. In biofloc system, bacteria could product exogenous digestive enzymes, increasing the digestibility of foods and improving the growth of shrimp (Panigrahi, Esakkiraj, Jayashree, Saranya, Das, & Sundaram, 2019; Wang, Pan, Zhang, Xu, Zhao, & Mei, 2016; Xu, Pan, Sun, & Huang, 2013). Additionally, inferring functions of bacterial community relating to biosynthesis of immunostimulants (LPS and peptidoglycan) and antibiotics (ansamycins, streptomycin and vancomycins) were observed in the current study, which would be helpful to maintain the high survival rate of PL. Exposure to stimulation of immunostimulants could reinforce the immune system of shrimp (Li & Xiang, 2013; Panigrahi, Sundaram, Saranya, Swain, Dash, & Dayal, 2019), and functions of biosynthesis of antibiotics might partially explain why bacterial pathogens do not have the same virulence and are inhibited in biofloc systems (Ekasari, Azhar, Surawidjaja, Nuryati, Schryver, & Bossier, 2014; Panigrahi, Sundaram, Saranya, Kumar, Dayal, Saraswathy, … & Gopal, 2019). In the current study, proportion of family *Vibrionaceae*, the most important opportunistic pathogenic bacteria groups for *L. vannamei* (Gomez-Gil, Tron-Mayén, Roque, Turnbull, Inglis, & Guerra-Flores, 1998; Kita-Tsukamoto, Oyaizu, Nanba, & Simidu, 1993), was only 0.003%.

### 4.5 Influence of bacterial community on water quality

The levels of ammonia, nitrite and nitrate at 28 d were low in this study, indicating that weak nitrification in the system. Actually, Only two families belonging to autotrophic nitrifying bacteria, *Nitrospiraceae* and *Nitrosomonadaceae* (Sliekers, Derwort, Gomez, Strous, Kuenen, & Jetten, 2002), were found in the bacterial community at 28 d of the present study, with a total proportion of 0.004%. Besides, although some heterotrophic bacteria with nitrification capacity were also found in the bacterial community of the present study, such as genera *Acinetobacter, Bacillus, Paracoccus* and *Pseudomonas* (Chen, Luo, Meng, & Tan, 2019; Liu, Li, Wei, Zhu, Han, Jin, … & Xie, 2019), their proportions were also very low. Correspondingly, the proportion of inferring functions of bacterial community relating to inorganic transformation were low (18% of the level-2 KEGG pathway nitrogen metabolism) in the present study. Conversely, conversion of ammonia to glutamate was the most important KO functions relating to the level-3 KEGG pathway of nitrogen metabolism in this study. Coincidently, total nitrogen and biofloc significantly accumulated at the end in this study, indicating massive formation of organic nitrogen or bacterial biomass. Heterotrophic bacteria could assimilate inorganic nitrogen to synthesize self-cellular protein, and as a result, massive bacterial biomass produced and biofloc formed (Ebeling, Timmons, & Bisogni, 2006). Impressively, the level of biofloc volume increased to 127.0±63.0 mL L^-1^ at the end of the experiment (28 d) in the present study, far higher than the acceptable value for shrimp (Avnimelech, 2015; Xu & Pan, 2012), meaning that removal treatment should be executed in next studies, but the effect of this treatment on bacterial community should be investigated as well.

It should be noted that in the present study, only the bacterial community at the end of experiment (28 d) was determined, leading to that only qualitative relationship between bacterial community and water parameters determined at the same time could be speculated, and that no tighter associations or quantitative correlations could be obtained. Thereby, interestingly and meaningfully, more treatments would be built up to get deeper insight of effects of bacterial community on growth performance and water quality in next studies.

## 5 CONCLUSION

The bacterial community at 28 d in the biofloc system rearing *L. vannamei* PL with a low salinity of 5.0‰ was highly diverse, and dominated by Proteobacteria, Bacteroidetes, Planctomycetes and Firmicutes. Those dominant species contributed to most of inferring functions of the bacterial community. Among the functions relating to nitrogen metabolism, 48.5% were involved in the conversion of ammonia to glutamate, but the proportion of those involved in transformation among ammonia, nitrite and nitrate was 18.0% in total, inferring higher protein-synthesis but lower inorganic nitrogen compounds-transformation capacities of the bacterial community, which were consistent with the practical levels of nitrogen compounds in water body at the same time (28 d), suggesting important roles of bacterial community in biofloc system.

## ACKNOWLEDGEMENTS

This work is financially supported by the development funds of the Chinese central government to guide local science and technology (2017CT5013), the Sci-Tech program of Hunan province (2016NK2132), the scientific research program of the Education Department of Hunan province (18B394), the open projects of Key Laboratory of Health Aquaculture and Product Processing in Dongting Lake Area of Hunan Province (2018KJ001) and Changde Research Center for Agricultural Biomacromolecule (2020AB08).

## AUTHOR CONTRIBUTIONS

Hai-Hong Huang: designed study and drafted the paper. Ting Luo: performed laboratory analysis. Yan-Ju Lei: analyzed the data. Wei-Qi Kuang: performed metagenomics analysis. Wan-Sheng Zou: performed laboratory analysis. Pin-Hong Yang: involved in English edition.

## DATA AVAILABILITY STATEMENT

Data are available if required.

